# A semi-analytic elastic rod model of pediatric spinal deformity

**DOI:** 10.1101/2020.04.20.051987

**Authors:** Sunder Neelakantan, Prashant K. Purohit, Saba Pasha

## Abstract

The mechanism of the scoliotic curve development in healthy adolescents remains unknown in the field of orthopedic surgery. Variations in the sagittal curvature of the spine are believed to be a leading cause of scoliosis in this patient population. Here, we formulate the mechanics of S-shaped slender elastic rods as a model for pediatric spine under physiological loading. Secondarily, applying inverse mechanics to clinical data of the scoliotic spines, with characteristic 3D deformity, we determine the undeformed geometry of the spine before the induction of scoliosis. Our result successfully reproduces the clinical data of the deformed spine under varying loads confirming that the pre-scoliotic sagittal curvature of the spine impacts the 3D loading that leads to scoliosis.

## 1. Introduction

The etiology of the adolescent idiopathic scoliosis (AIS) remains largely unknown [27, 8]. Several hypotheses have been developed to explain the patho-mechanism of AIS development [20, 16, 12, 5, 9]. Among these hypotheses, the upright alignment of the spine in humans, which impacts the mechanical loading of the spine, is believed to be an important factor in induction of scoliosis [6, 20]. The shape of the sagittal curvature of the spine, prior to initiation of spine deformity development, has been shown to be different between the scoliotic and non-scoliotic age, sex matched cohorts [26]. The shape of the sagittal curvature of the spine in pre-scoliotic patients was believed to make the spine rotationally unstable and lead to scoliosis [20, 7]. However, as the pre-scoliotic data on the sagittal profile of this patient population is scarce, an analysis that can determine the physiologically acceptable undeformed shapes of the sagittal spine that can lead to scoliotic-like deformation can be valuable for early clinical diagnosis of the curves. Identifying the characteristics of the spinal sagittal curvatures that are prone to scoliotic curve development under a general set of loadings can be used as a risk stratification tool for early diagnosis of the disease.

Our strategy to efficiently identify sagittal curvatures of the spine under loads is to model it as an elastic rod. The human spine has already been treated as an elastic object in finite element calculations that are routinely performed to compute its deformation under loads [20]. Since the spine is a slender structure whose length dimension is much longer than the cross-sectional dimensions we treat it as an elastic rod in this work. For simplicity, we assume that the rod has a circular cross-section and is inextensible. Both these assumptions are introduced for convenience and can be relaxed in a more general theory [1].

Since the material of the spine can be modeled as linearly elastic for small strains [11], our rod model for the spine has two equal bending moduli *K*_*b*_(*s*) and a twisting modulus *K*_*t*_(*s*) which vary as a function of position *s* along the centerline of the cross-section. The moduli are allowed to vary as a function of position because the cross-sectional dimensions of the spine as well as the material properties of the spine are different depending on the position. For example, the thoracic region has a higher modulus compared to the lumbar region due to attachment of the rib cage. Furthermore, we wanted to allow for the possibility that a scoliotic spine may have these properties varying in a different manner compared to a healthy one. We also assume that the stress-free configuration of our rod model for the spine is S-shaped as observed in a human upright standing position. Thus, our rod has a spontaneous curvature *κ*_0_(*s*) (which is inversely proportional to the radius of curvature) which is a function of position along the centerline. This stress-free curvature is assumed to have zero twist component for a healthy spine. The advantage of modeling the spine as a curved elastic rod under loads is that its deformed shapes can be computed by solving a few ordinary differential equations rather than performing a full finite element calculation which requires many input variables. We show in this paper that our model, although simple, can reproduce many important features in the deformation of spines that have been observed clinically and studied in finite element calculations.

This study aims to determine the deformation patterns of various S-shaped elastic rods as are observed in sagittal curvature of the scoliotic spines. It also aims to match the simulated deformation of such a rod model to the clinical data by altering the mechanical loading and mechanical properties of the rods. We hypothesized that an elastic rod under bending and torsional moments deforms in a similar manner as a pediatric spine with scoliosis. Also, the rod’s deformation and clinical data of the corresponding curved spines can produce similar deformations by altering the mechanical loading of the rods.

## 2. Methods

### 2.1. Clinical data and data preparation

The deformed sagittal curvatures of the scoliotic spine were determined from a previous classification study of 103 scoliotic patients [22]. In brief, these 3D curves were generated by post-processing of the clinical radiographs [24]. A hierarchical clustering determined the subsets of the patients in this cohort with five significantly different 3D spinal curves. The five cases are presented below in fig.(1). It was shown that these five subtypes can be divided into two groups based on the top-down view (X-Y) of the 3D curves: cases 1, 3 and 5 have lemniscate shaped (figure 8-shaped) X-Y view and cases 2 and 4 have loop shaped X-Y view. We used these two patterns (loop and lemniscate) as the basis of our study to describe, using forward and backward mechanics, the parameters causing an undeformed spine to develop either of these deformation patterns. These views are also known as sagittal view (Y-Z view), frontal view (X-Z view), and axial view (X-Y view).

**Figure 1:**
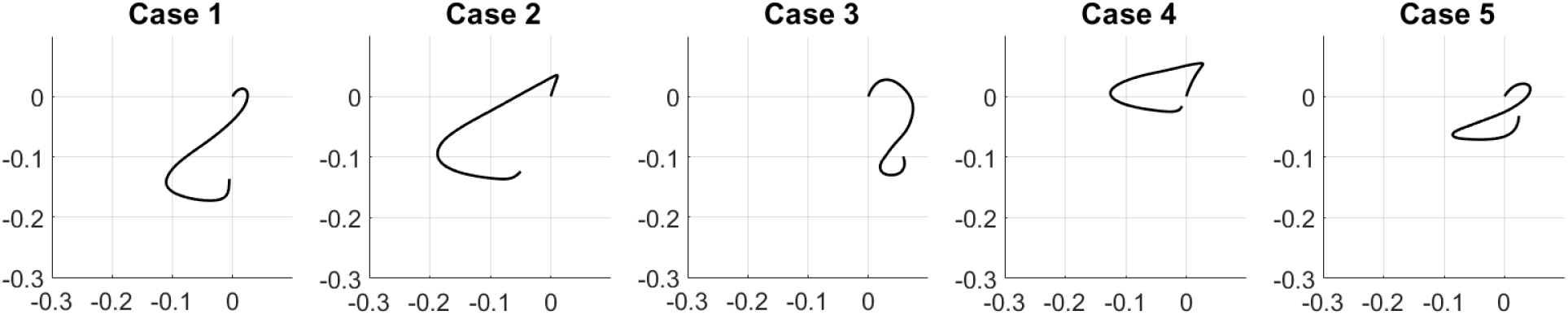
The axial view of the 5 cases of scoliosis

### 2.2. Elastic Rod Model

Let us consider the undeformed spine as an S-shaped elastic rod. The rod is assumed inextensible. The sagittal plane of the spine is co-incident with the *y* − *z* plane of the lab frame. The undeformed rod is assumed to have no out-of-plane curvature with respect to the sagittal plane. The origin of the lab coordinate system [**e**_*x*_ **e**_*y*_ **e**_*z*_] is placed at the bottom end of the spine *s* = 0 where *s* is an arc length coordinate along the center line of the spine. The position of a point *s* in the deformed configuration is **r**(*s*) = *x*(*s*)**e**_*x*_ + *y*(*s*)**e**_*y*_ + *z*(*s*)**e**_*z*_.

Now we define the Frenet frame for the rod, given by the triad 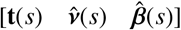. Here 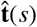 is the tangent vector and is given by the equation 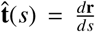. It is a unit vector because the rod is assumed inextinsible. The tangent is given by 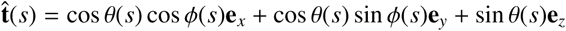. Here *θ* is the polar angle measured from the *x* − *y* plane and *ϕ* is the azimuthal angle used in conventional spherical polar co-ordinates. The conventions and variables are described in fig.(2). 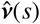 and 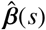 are the normal and binormal vectors, respectively. They are related through the Frenet-Serret equations [19].

**Figure 2:**
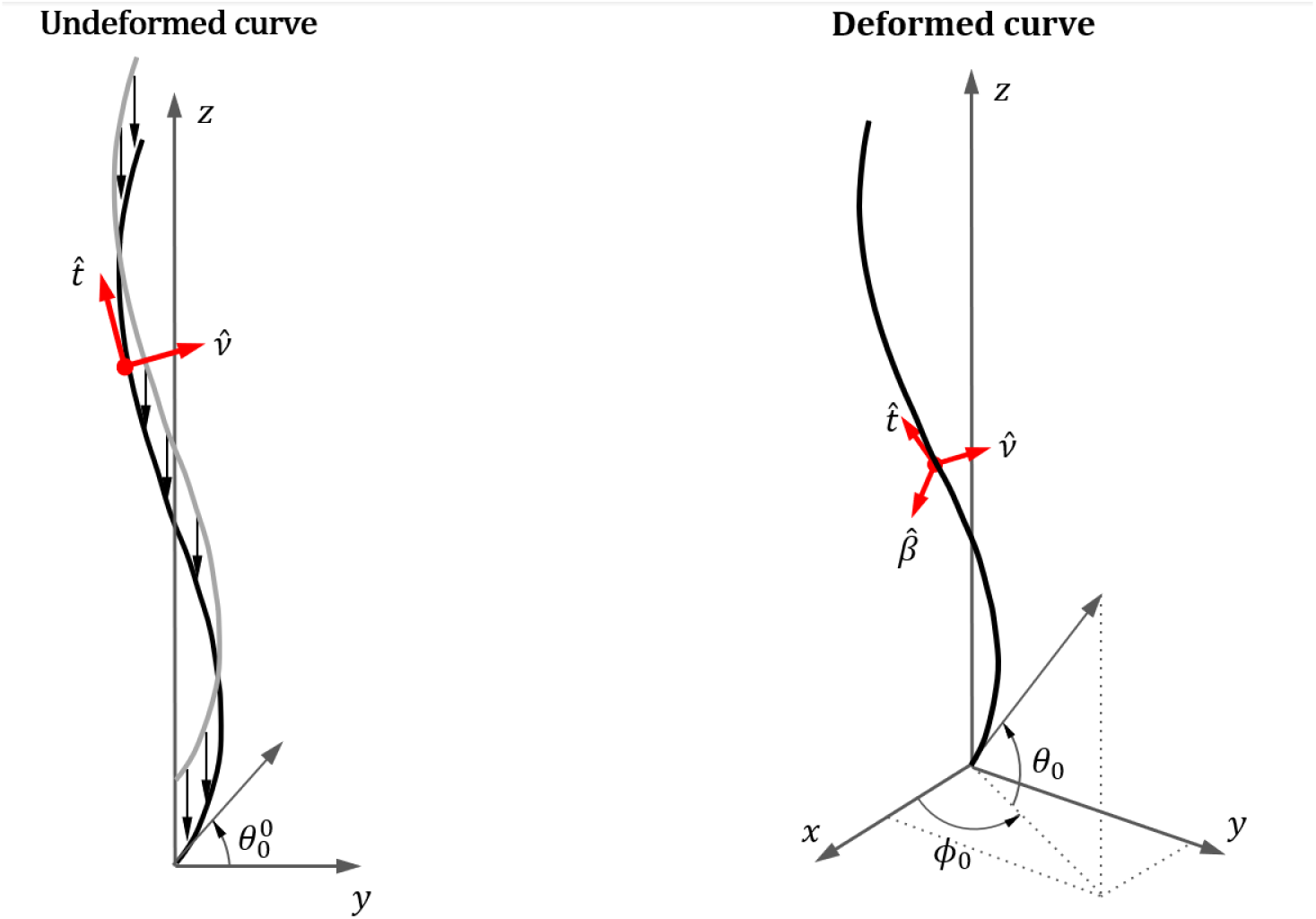
The centerline of the rod a before and after deformation. The [**x y z**] system of co-oridnates represent the lab frame. The 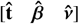 system represents the Frenet-Serret frame for the deformed and undeformed configurations

The curvature and torsion of the curve describing the center-line of the spine are *κ*(*s*) and *τ*(*s*), respectively, and are given by:

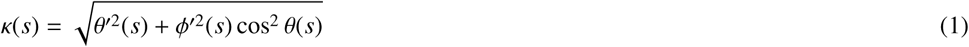

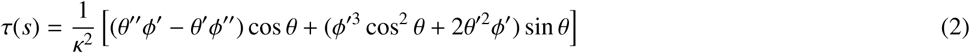

#### 2.2.1. Forward mechanics

This section will deal with the mechanics of the rod. We will present an analytical model to compute the deformed geometry for a given S-shaped rod. The balance of linear momentum [1, 2] gives

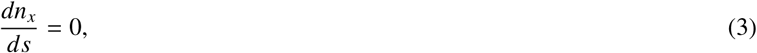

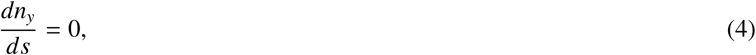

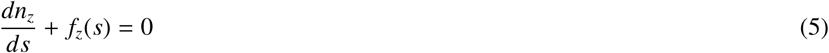

where **n**(*s*) = [*n*_*x*_(*s*) *n*_*y*_(*s*) *n*_*z*_(*s*)] is the force in the rod and the distributed load **f** = *f*_*z*_(*s*)**e**_*z*_ is directed only along the **e**_*z*_ direction due to gravity. It follows immediately from the above that

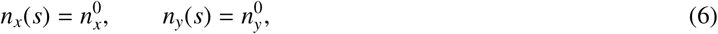

where 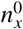 and 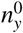 are constants that will be determined by the boundary conditions. Since there are no forces applied on the spine along **e**_*x*_ and **e**_*y*_ directions, then global equilibrium shows that 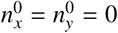.

For the balance of angular momentum, the moment in the rod **m**(*s*) is represented in the Frenet frame and given by 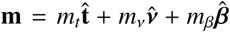 at any point and **l** is a body moment per unit length. We will take **l** = 0 in this work. A simple constitutive relation for the moment **m** is

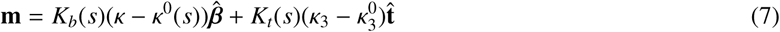

where *κ*_3_ is the twist rate and *K*_*b*_ and *K*_*t*_ are the bending and twisting moduli of the spine. We assume that *K*_*b*_ and *K*_*t*_ are functions of *s* since there is variation in spine stiffness in a scoliotic spine and also because the material properties of the spine may vary as a function of position. 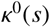 and 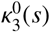 are the values of the curvatures in the stress free state, respectively. We assume 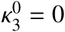. This constitutive law is of the form 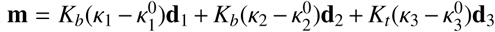 where [**d**_1_(*s*) **d**_2_(*s*) **d**_3_(*s*)] is a material frame that convects with the arc-length coordinate *s* along the center-line of the cross-section^1^. Now, the derivative of the moment becomes

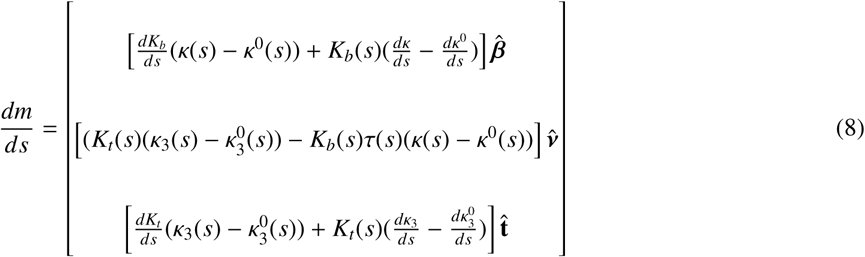

Hence the conservation of angular momentum boils down to [2, 1]

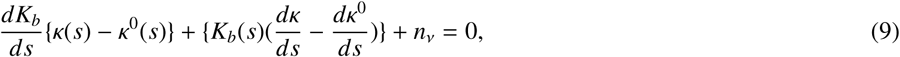

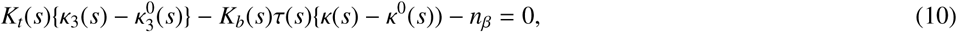

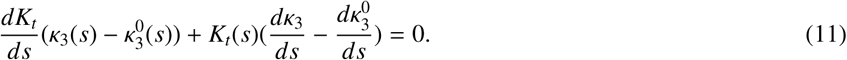

We define

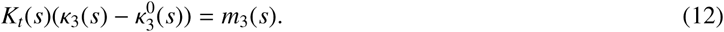

Then, eqn. (11) shows that

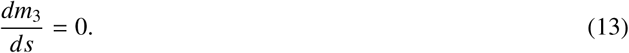

Hence, the twisting moment in our rod model is constant along the arc-length. We can also write the conservation of angular momentum in the lab frame as

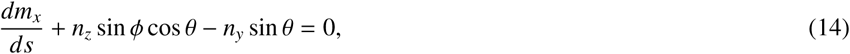

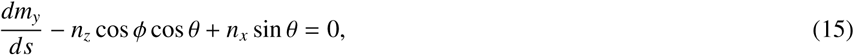

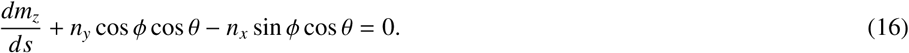

If *n*_*x*_ = *n*_*y*_ = 0, as we concluded from the balance of linear momentum, then the third equation above gives *m*_*z*_ = *T*, a constant that is determined by a torque boundary condition applied at *s* = 0.

Finally, the moment expression is given by:

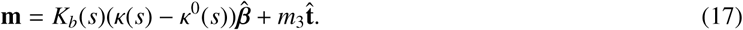

We express this moment in the lab frame as:

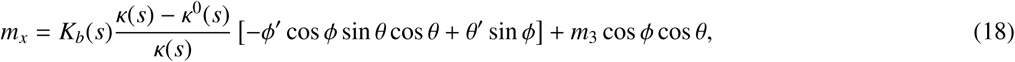

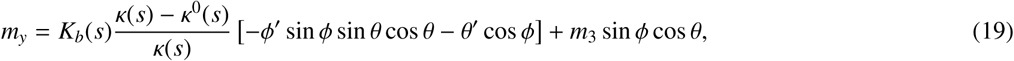

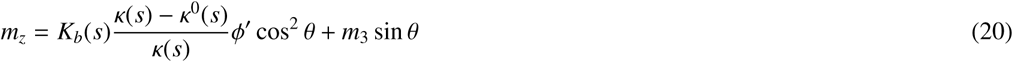

To better understand the effects of moments, we will describe the effect of a constant moment along a single direction on the spine (S-shaped rod). A moment along 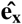 axis causes the spine to straighten the lumbar curvature and increase the thoracic curvature (leading to kyphosis). A moment along 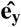 axis would cause a person to tilt side-ways. A moment along 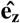 would cause the spine to twist to resemble a helix. Going back to the equations for *m*_*x*_, *m*_*y*_, *m*_*z*_ above, *K*_*b*_(*s*), *κ*^0^(*s*) and *κ*(*s*) can be eliminated to give:

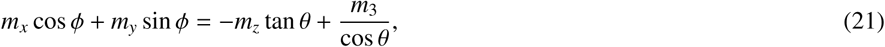

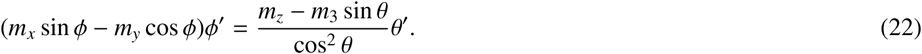

Eqn. (21) can then be solved to get

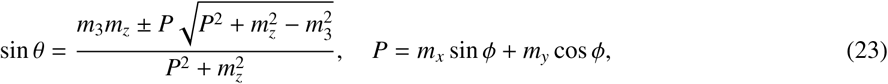

where the solution branch can be determined from the initial value of *θ*(*s*). We can find an expression for *ϕ′*(*s*) using eqn. (22) and eqn. (2.2) to get

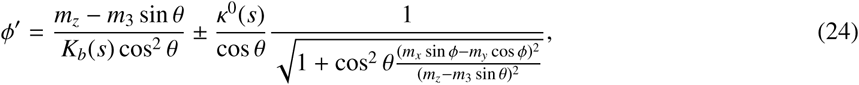

where the ± sign is dependent on the sign of *ϕ′*(*s*).

Finally, the analytic model of the spine is given by the following system of differential-algebraic equations.

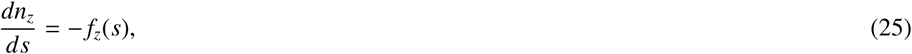

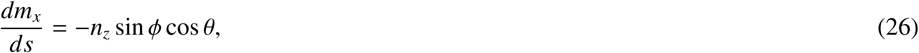

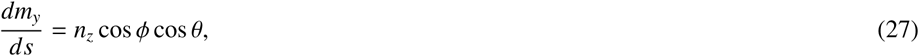

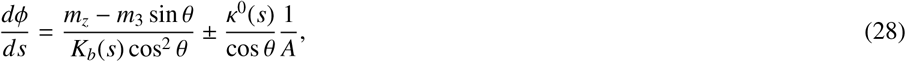

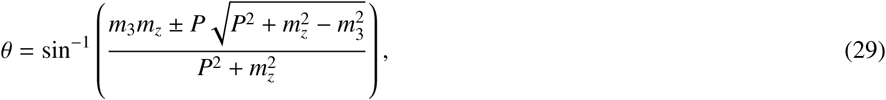

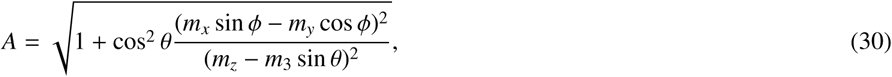

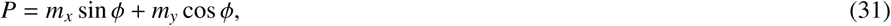

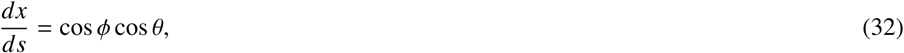

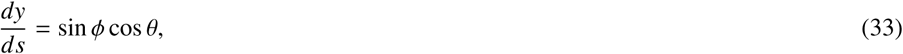

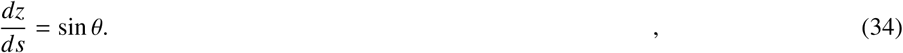

In the above the body force *f*_*z*_(*s*) can be found from a generalized load distribution along the upper body as given by Pasha et. al[21].

#### 2.2.2. Inverse mechanics

In this section, we will present a method to extract the properties of the undeformed rod and the forces acting on it, given the deformed geometry. The deformed geometry has been obtained from clinical data of patients suffering from scoliosis and is given as a set of position vectors of points along the deformed spine[22]. We interpolate the data to obtain a larger array of points which are spaced uniformly and closer together than the clinical data. The size of the larger array is *N* (which can be set) and each point be given by *p*_*i*_ = [*x*_*i*_ *y*_*i*_ *z*], where *i* goes from 1 to *N*. Then, the length of the spine can be computed by

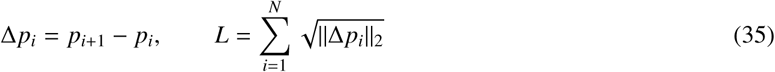

Consider *s* as an array of N points which can be defined by 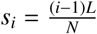. Hence, we can compute *θ* and *ϕ* by

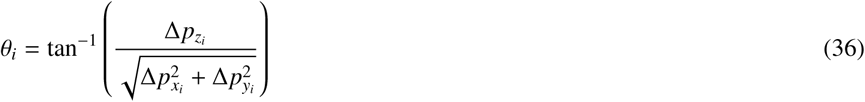

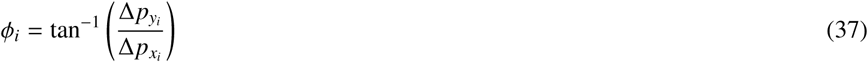

We can then use *θ*_*i*_ and *ϕ*_*i*_ to compute their derivatives with respect to *s* i.e 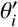, and 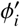; we use these to compute the deformed curvature *K*_*i*_. Then, we use *f*_*z*_(*s*) to determine the moments.

#### 2.2.3. Moment calculation from generalized body force

Recall the governing system of differential-algebraic equations above, specifically eq. (25), (26) and (27). We can compute the values of *m*_*x*_(*s*) and *m*_*y*_(*s*) since we have values of θ and ϕ. However, we do not know the initial values (at *s* = 0) of the moments. To find *m*_*x*_(0), *m*_*y*_(0), *m*_*z*_ and *m*_3_, we minimize the least squares error between eq. (29) and the clinical data. This is given by

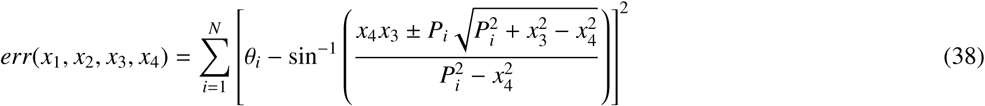

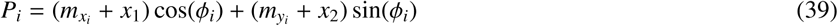

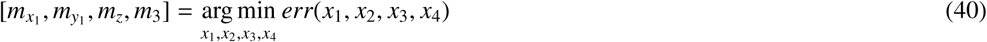

Hence, we offset the *m*_*x*_ and *m*_*y*_ values by *m*_*x*_1 and *m*_*y*_1 respectively. Now we use the moment values to determine the *K*_*b*_(*s*) and *κ*^0^(*s*). We use eq. (20) to define an error term based on the least squares method to determine *K*_*b*_(*s*) and *κ*^0^(*s*).

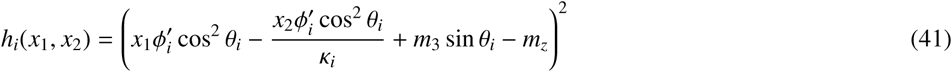

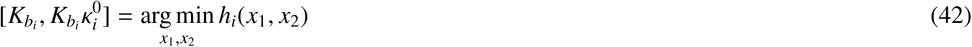

subject to the condition 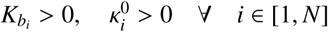.

After verifying the values from the minimization, we compute the spinal geometry *prior* to twisting using a special case of the ODEs with *ϕ*(*s*) = *π*/2, implying that *ϕ′* = 0. For this case, we assume that the spine is a planar rod being deformed by *f*_*z*_(*s*) alone. This leads to out of plane moments going to 0. i.e *m*_*y*_(*s*) = 0, *m*_*z*_ = 0, *m*_3_ = 0 with only *m*_*x*_(*s*) being the non-zero moment. We get the geometry by solving the following differential equations.

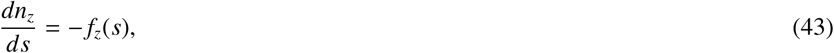

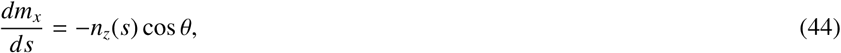

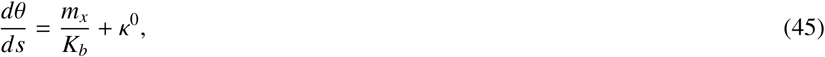

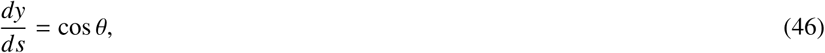

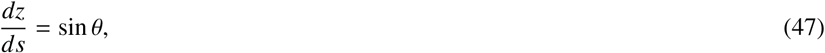

with the initial conditions being

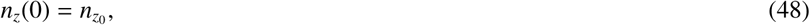

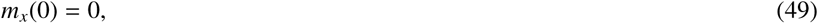

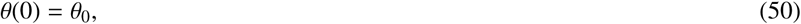

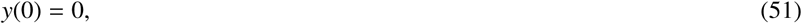

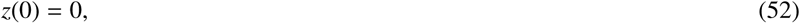

where 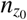 is the weight of the upper body and *θ*_0_ is the base angle of the spine in the sagittal plane at *s* = 0.

We presented these curves in fig.(7). We chose *κ*^0^(*s*) to ensure that the rod remains upright. We constrain the horizontal displacement of the top of the rod to remain under 15% of the vertical displacement. We set this bound to ensure that the head remains roughly over mid-line of the body in the sagittal plane

Here, we define a few terms for ease of understanding. We define *S* as the point of inflection of the rod, i.e., where the curvature approaches 0. *K*_*P*_ is average of the curvature values of the part of rod occupying *s* < *S*, i.e 1/*K*_*P*_ is the radius of curvature of the lumbar region of the spine. *K*_*N*_ is the average of the curvature of the part of the rod occupying *s* > *S*, i.e 1/*K*_*N*_ is the radius of curvature of the thoracic region of the spine. We present these values in table1.

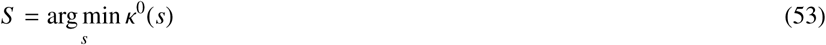

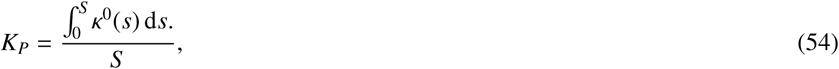

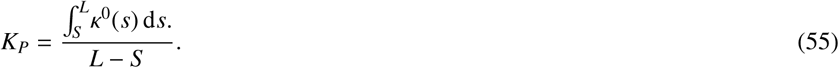

**Table 1:**
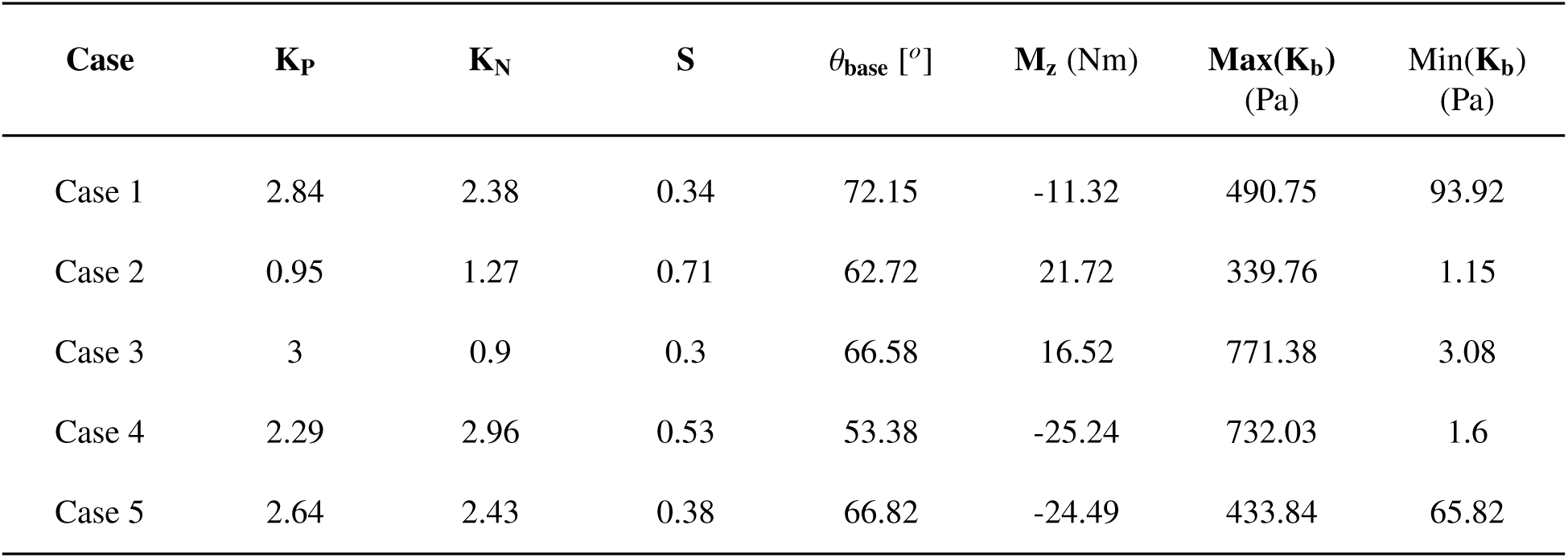
Table of average curvature values of the thoracic and lumbar regions

| **Case** | **$K_P$** | **$K_N$** | **S** | **$\theta_{\text{base}}$ [$^\circ$]** | **$M_z$ (Nm)** | **Max($K_b$)**<br>**(Pa)** | **Min($K_b$)**<br>**(Pa)** |
| --- | --- | --- | --- | --- | --- | --- | --- |
| Case 1 | 2.84 | 2.38 | 0.34 | 72.15 | -11.32 | 490.75 | 93.92 |
| Case 2 | 0.95 | 1.27 | 0.71 | 62.72 | 21.72 | 339.76 | 1.15 |
| Case 3 | 3 | 0.9 | 0.3 | 66.58 | 16.52 | 771.38 | 3.08 |
| Case 4 | 2.29 | 2.96 | 0.53 | 53.38 | -25.24 | 732.03 | 1.6 |
| Case 5 | 2.64 | 2.43 | 0.38 | 66.82 | -24.49 | 433.84 | 65.82 |

We can see the effects of the moments better in the Frenet frame of the rods. We compute the Frenet frame 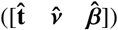 for the curve prior to the twist (presented in fig.(7)) as.

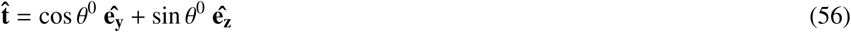

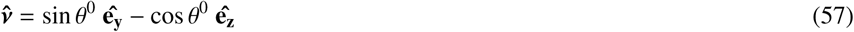

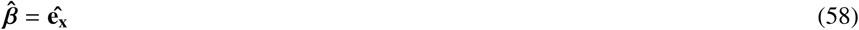

where *θ*^0^ is defined in fig.(2). We then decompose the moments along this frame and plot *m*_*nu*_(*s*) on the pre-twist curve in fig.(8). Now *m*_*β*_(*s*) is responsible for the shape of the spine in the sagittal view while *m*_*ν*_(*s*) is responsible for the shape of the spine in the frontal view. We will present an explanation using case 5 as an example in the discussion section.

## 3. Results

We applied the inverse mechanics method to the five clinically derived cases to determine the stiffness, undeformed curvature and moments. We validate the model by using the parameters to solve the system ODEs(eq.32, 33, 34). We present a comparison between the clinical data and the model in the axial view in fig.(3) and the sagittal view in fig.(4) and the frontal view in fig.(5). These figures shows that the model is in good agreement with clinical data. Plots of *m*_*x*_(*s*) and *m*_*y*_(*s*) are presented in fig.(6). The moment and stiffness values of the spine are presented in Table(1). We used these model parameters to predict the curvature using eq. (42). The shapes of the spine prior to the twisting effects for each of the cases in fig.(1) are presented in fig.(7).

**Figure 3:**
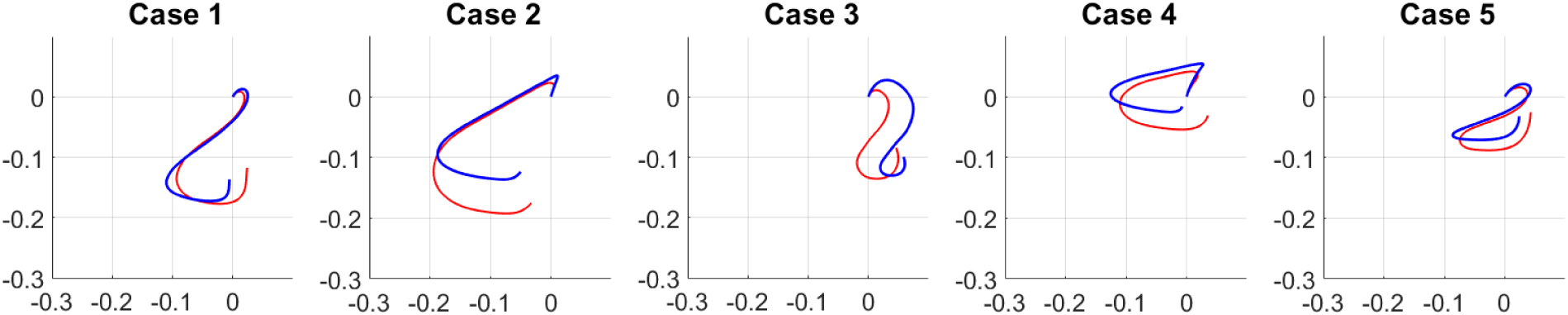
Comparison between the clinical data and the computational model in the axial view. The red curves are the results of the computational model. The blue curves are clinical data from [22].

**Figure 4:**
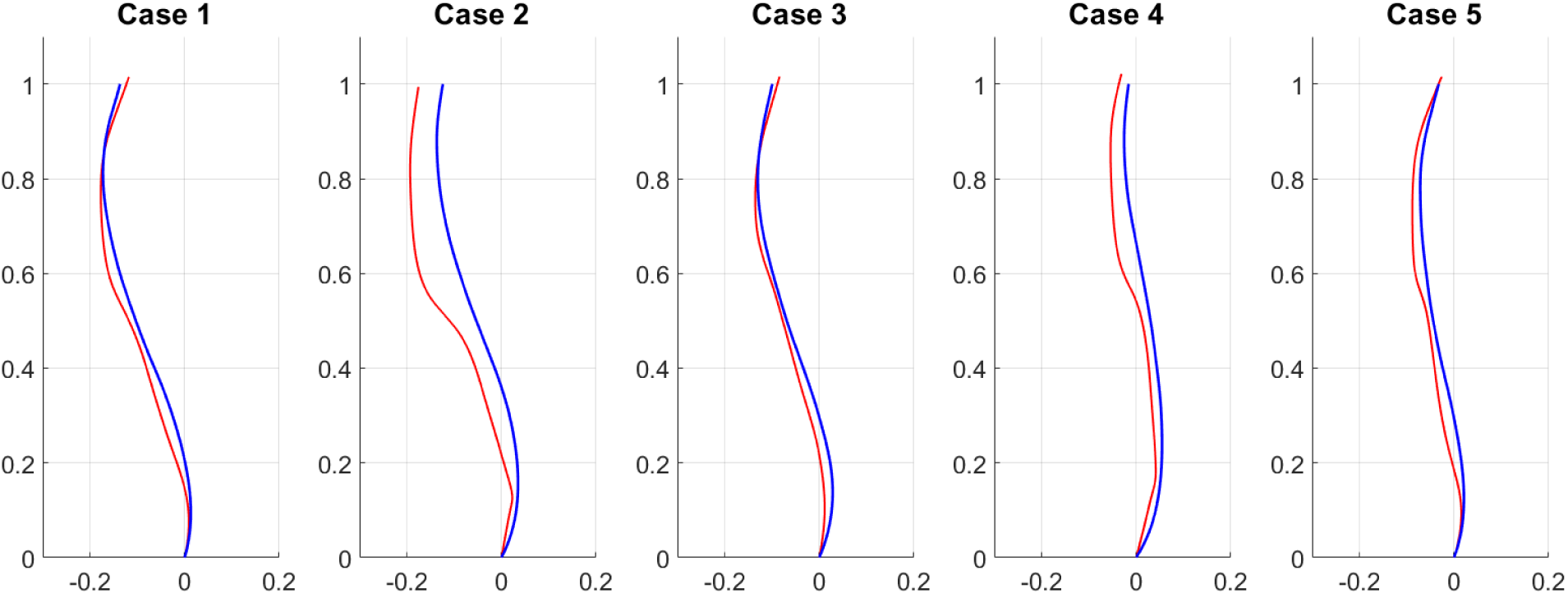
Comparison between the clinical data and the computational model in the sagittal view. The red curves are the results of the computational model. The blue curves are clinical data from [22].

**Figure 5:**
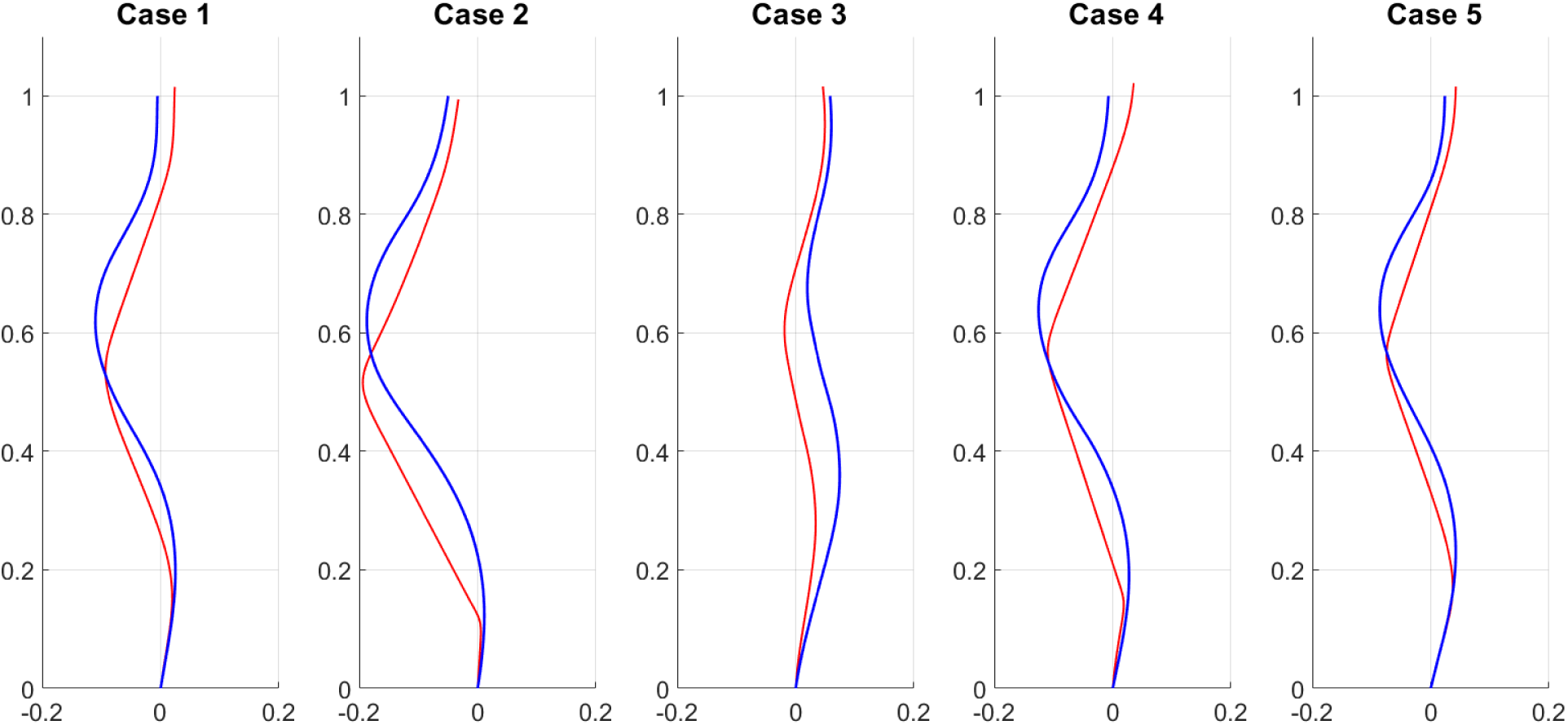
Comparison between the clinical data and the computational model in the frontal view. The red curves are the results of the computational model. The blue curves are clinical data from [22].

**Figure 6:**
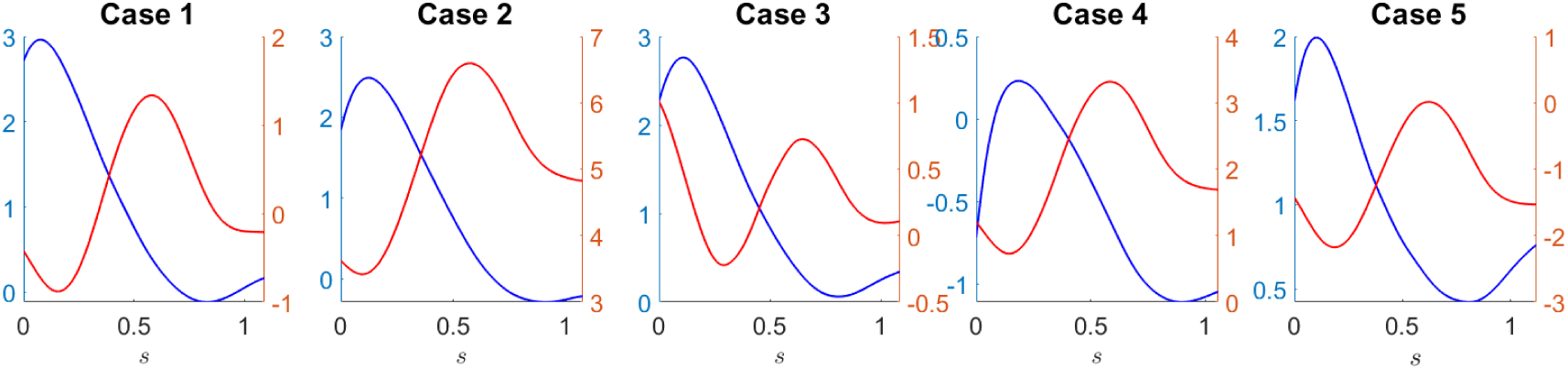
Trends of *m*_*x*_(*s*) and *m*_*y*_(*s*) for the 5 cases. The blue curves are *m*_*x*_(*s*) correspond to the left y-axis. The red curves are *m*_*y*_(*s*) correspond to the right y-axis.

**Figure 7:**
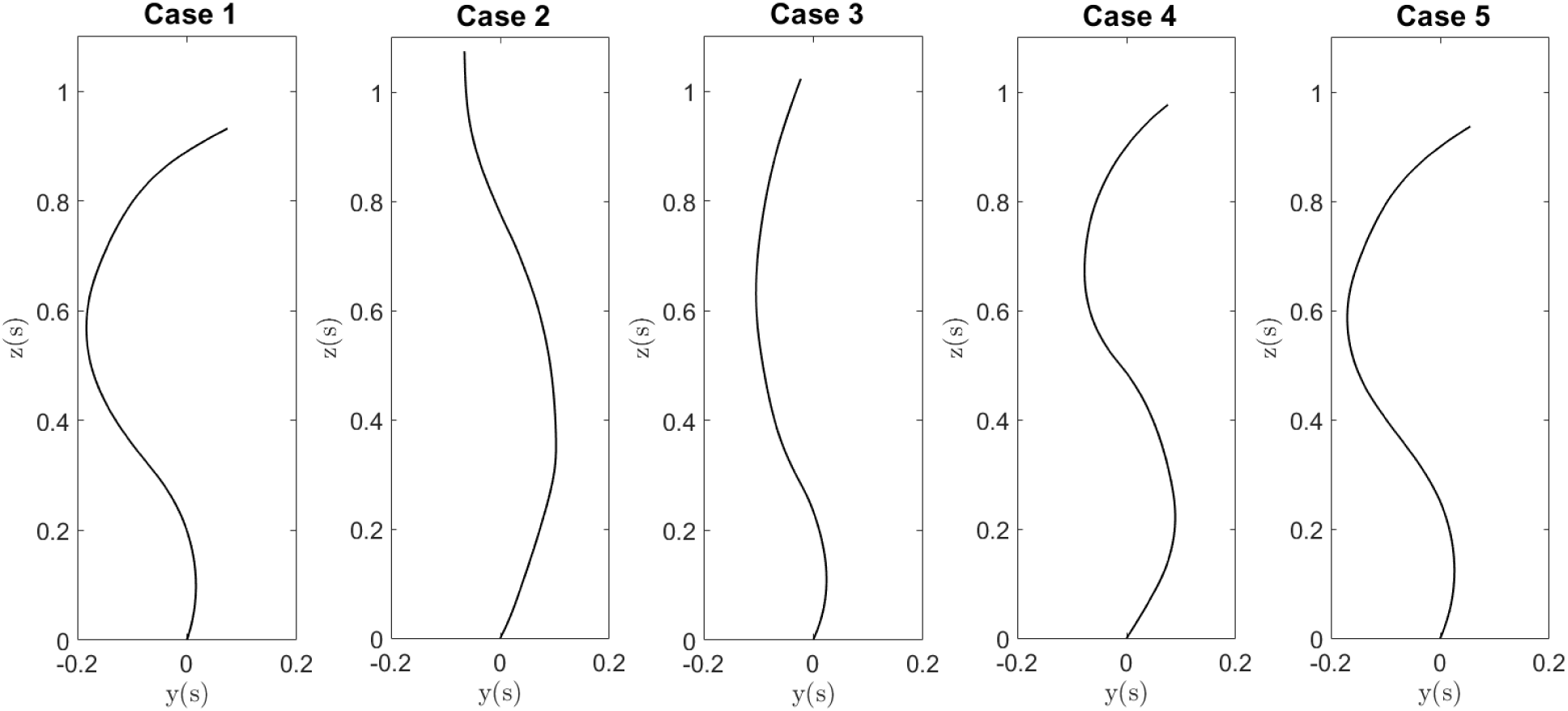
The geometry of the spine before twist in the sagittal view

We present the curve prior to twist and the *m*_*ν*_(*s*) component of the moment decomposed along the Frenet frame for the curves presented in fig.(8) to better understand the physical significance of the moments and the effect they have on the deformed shape in the axial view.

**Figure 8:**
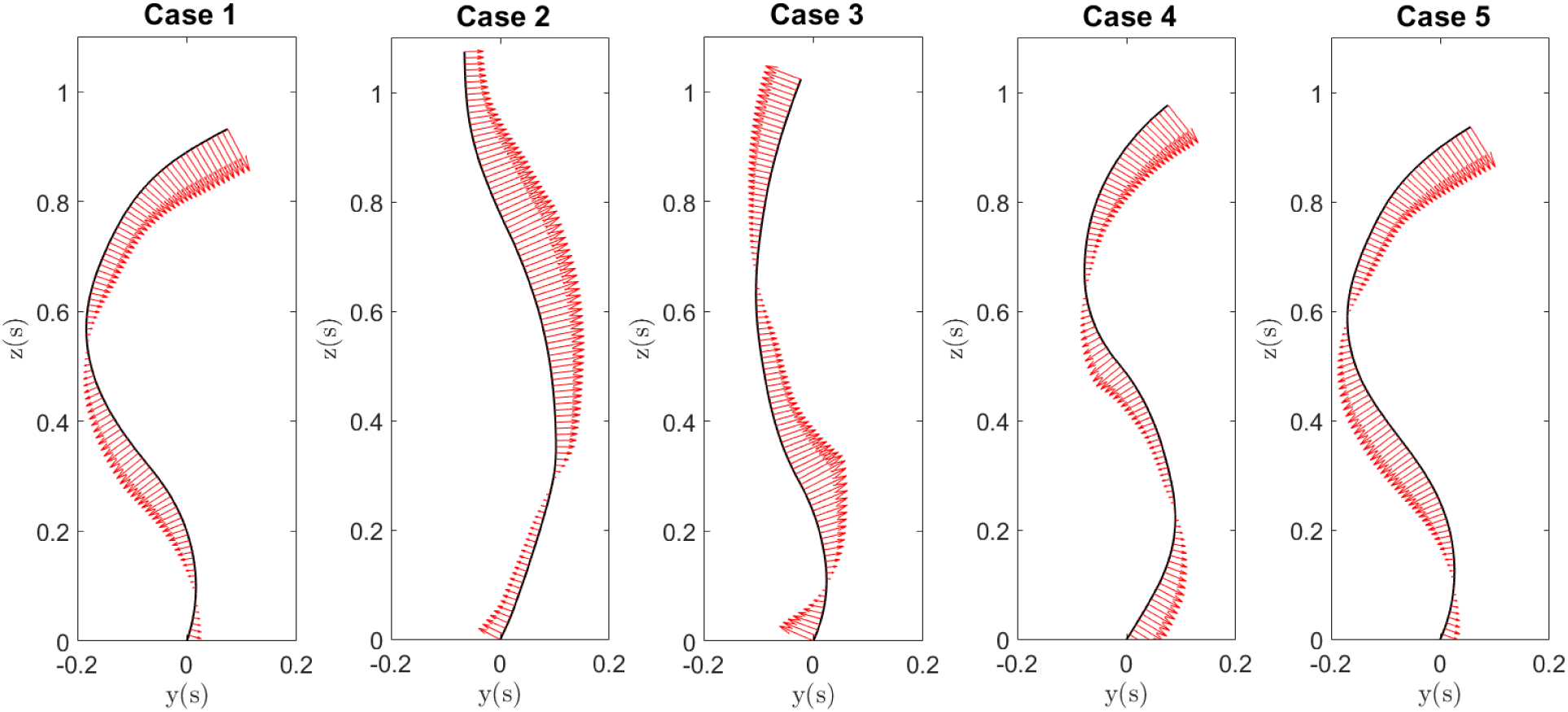
*m*_*nu*_(*s*) plotted on the pre-twist curve. Vectors point away from the curve.

We compute the average curvatures for the 2 parts of the S-shaped rod and the point of inflection and present them in the table 1. The *m*_*z*_ experienced in each case along with the variation in stiffness is also presented in the table 1 We also present a representative stiffness curve for this model in fig.(9) (Taken from case 5).

**Figure 9:**
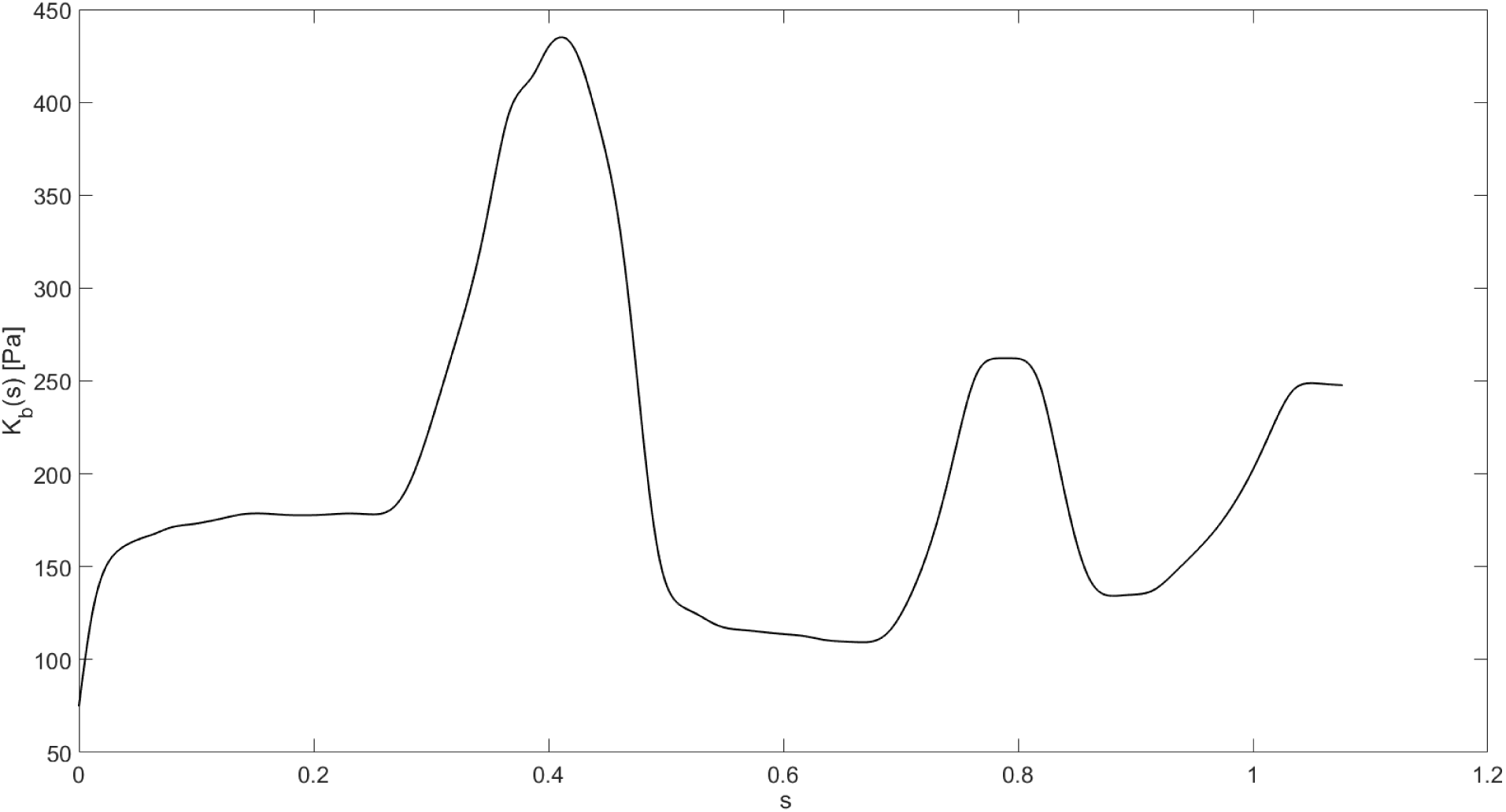
A representative *K*_*b*_(*s*) curve. This is produced from Case 5.

## 4. Discussion

The enigmatic spinal deformation in adolescents has been studied for centuries. Here we use an analytical model of a curved elastic rod to study the fundamentals of the 3D deformations as it relates to the curve development in scoliosis. We use the clinical subgroups of scoliotic patients with a thoracic curve and apply inverse mechanics to determine the undeformed shape of the spine. In this analysis, by *untwisting* the S shaped rod under gravity, we determine the shape of the spine before the induction of scoliosis. Our results show the characteristic of the S-shaped curvatures i.e. sagittal profile of the spine, was preserved after untwisting the curve, meaning that the spine with a loop shaped projection (Cases 2 and 4) have a larger lordosis than kyphosis with an inflection point above the center of curve, whereas spines with a lemniscate axial projection (Cases 1, 3 and 5) have a larger kyphosis than lordosis with an inflection point below the center of the curve. We compare the deformed S-shaped curves under gravity and torsion to the clinical data and find acceptable agreement between the simulations and clinically reported deformity patterns as seen in fig.(3) and fig.(4).

The sagittal curvature of the spine is believed to have an important role in induction of scoliosis [15, 20]. However, the data on the sagittal curvature of the spine prior to curve development is scarce. In a previous study, Pasha et. al showed, using finite element modeling, that an S-shaped elastic rod under bending and torsion can deforms in loop or lemniscate shaped in the axial projections only as a function of the curve geometry as seen in fig.(1). When those curve geometries were compared to the clinical data, it was observed that sagittal curvature of the scoliotic patients also related to the axial projection of the curve, in the same manner as an elastic rod. However, as that study used the sagittal profile of the scoliotic curves, it could not be shown what characteristics of the sagittal plane determined the deformity patterns of the spine in scoliosis. Our analysis in this paper uses defined scoliotic curve types and applies inverse mechanics analysis to determine the pre-scoliotic shape of the spine. These shapes are then shown to produce the 3D scoliotic deformity, under physiologically acceptable conditions, that matches the clinical data. The current analysis shows that the moments that cause the off-plane curve deformation can be formulated as a function of the curves sagittal parameters as seen in fig.(8). This study explains how the initial sagittal curvature of the spine impacts the mechanical loading of the spine, which in turn leads to the scoliotic like deformation. Understanding the fundamentals of the spinal loading and resultant deformities is the first step in developing clinical methods that prevent or reverse the deformity development.

While several hypothesis have been developed to explain the spinal deformity in AIS, the use of analytical models remains unexplored. The only other existing analytical model of the spine for scoliosis, explains this deformity only in one plane (frontal plane). These models solve the spine as a 2D straight rod thus ignoring the curvature of the spine in the sagittal plane. The deformity under gravity then was explained as 2D buckling of the rod. The deformation modes in the frontal plane were used to explain variations in the curve types in scoliotic patients[17, 3, 18]. But, AIS as we know it, is not 2D buckling. The curve deforms in 3D gradually. Our elastic rod model, as shown in this study, agrees with the characteristics of spinal deformity development in scoliosis as it incorporates the variation in the sagittal curvature. Such a deformation may be reversible if all deformations are elastic.

In the current model, in addition to gravity we used an axial moment to obtain the scoliotic shapes. This moment in the system was originally meant to break the symmetry of the system that would be otherwise only under gravity loads and thus would not deform in 3D. However, this torsion can be physiologically justified by the trunk mass asymmetry [10, 15, 13]. There is no evidence that this moment (torsion) is larger in scoliotic patients than in non-scoliotic patients or whether this moment varies between different curve types as a result of differences in the kyphosis and the chest volume. This merits investigation to further personalize the model.

We used several filtering criteria as the solution set to the inverse problem is not unique. We eliminated solutions that are physiologically not reasonable. We limited the maximum possible *m*_*z*_ and *K*_*b*_(*s*). We also ensured that the characteristic of the curve in the sagittal view was preserved. If one or more of these limiting conditions were not met, we repeated the optimization procedure with different initial values. This shows us that while the shape of the curve in the sagittal view is important in the induction of scoliosis, a unique curve cannot be produced given a deformed shape. We can determine a set of possible solutions with reasonable physiological parameters for a given deformed shape and can present a unique solution given more clinical data.

The spikes in *K*_*b*_(*s*) and *κ*^0^(*s*) are filtered out (to ensure that functions and their derivatives do not have discontinuities). the comparison between the clinical data and the solution from the model is presented in the results section. The solutions to eq. (40) and eq. (41) are not unique. The minimization function finds the local minima of the function in the neighborhood of the starting point. We select solutions by changing the start point. We selected the solutions based on physiological limitations, which we explain in the discussion section.

We limit our *K*_*b*_(*s*) values to 1000Pa based on a previous finite element simulations where they modeled the spine as a rod with a circular cross-section[20]. We present *K*_*b*_(*s*) in the supporting information and present the maximum and minimum values for the 5 cases in Table 2. The *K*_*b*_(*s*) (A representative curve is presented in fig.(9)) curve contains 3 distinct peaks. The peaks in *K*_*b*_(*s*) correspond to the peaks in the load function *f*_*z*_(*s*). As we simplified the load along the spine as point load with spikes to present the weight of the head and arms the Kb showed max value in the same points. In reality, it is expected as the loads are distributed more gradually the kb show a more smooth transition from region-to-region.

We also presented the corresponding *m*_*x*_(*s*) and *m*_*y*_(*s*) values in fig.(6). We can see that *m*_*x*_(*s*) are always positive for the lemniscate shapes while they change signs for the loop shapes. Looking at the *m*_*y*_(*s*) values, we see that *m*_*y*_(*s*) values always stay positive for the loop shape cases. This can be used as a criterion to determine whether loop or lemniscate shapes will develop. We also present plot of *m*_*ν*_(*s*) vectors plotted along the undeformed curve to understand the deformation in the X-Z plane. We do not consider the effects of *m*_*t*_(*s*) and *m*_*β*_(*s*) as they are responsible for twist and Y-Z plane deformations respectively. We can see in case 5 in fig.(8) that *m*_*n*_*u*(*s*) points towards the right -front of spineat the bottom and top regions and towards the left-backward- in the middle region and zero at the apices, but for cases 2 and 3 this relationship is reversed since the *m*_*z*_ was in opposite direction. This direction also relates to the θ which is less than 90◦ at bottom and top and exceeding 90◦ in the middle and 90◦ at the apices. We assume that the base is fixed, hence due to the moment, the curve would deflect towards the +ve x-axis in the bottom and top regions. The curve would deflect towards the −ve x-axis in the middle region. This is exactly what we see in fig.(5) for case 5. Hence we can predict the frontal view after we look at *m*_*ν*_(*s*) and since we know that the sagittal curve characteristic is roughly preserved, we can predict the shape of the curve in the X-Y projection.

The current model has several limitations that were required for constructing the analytical solution. First, the rod was considered to be inextensible. The changes in the sagittal alignment of the spine during the course of scoliotic development impacts the disc morphology as a function of mechanical loading and changes the curvature of the spine before any bony deformation occurs [23, 4, 25]. This mechanism may extend sections of the spine and contract the adjacent parts. Second, we considered the base angle as the tangent to the curve at the lowest vertebral level. While in reality the alignment of sacrum, which can be aligned independent from the shape of the spine at the lowest vertebral level, plays an important role in regulating the spinal alignment over the femoral heads and transferring the force between the spine and lower extremities [21, 14]. Considering the position of the spine over the sacrum may have required additional coordinate system transfer particularly in cases with large disc angulation above sacrum.

Despite the limitations mentioned above, our model delivers results which are in good agreement with clinical data. The model uses only five variables that can be easily captured in a clinical set up for patient assessment. This model is advantageous compared to the finite element models as it can save significant computational costs and time to achieve solutions of similar accuracy. We can calculate the forces and material properties using the **x, y, z** co-ordinates and the body force acting on the spine which are easily measurable.

## Acknowledgement

SN and PKP acknowledge partial support through an NSF grant NSF CMMI 1662101.

## Footnotes

1 It assumes that the stress-free curvature of the spine is aligned along the bi-normal vector *β*(*s*). This is certainly true for planar deformations of a healthy spine for which *x*(*s*) = 0 for all *s*. We show later that it also gives good results for full 3D deformations even though it is not the most general constitutive law.

